# Sex-biased computations underlying differential set shift performance in mice

**DOI:** 10.1101/2025.04.01.646712

**Authors:** Nic Glewwe, Evan Dastin-Van Rijn, Cathy S. Chen, Erin Giglio, Evan Knep, R. Becket Ebitz, Alik S. Widge, Nicola M. Grissom

**Affiliations:** University of Minnesota; Université de Montréal

**Author notes:** **to whom correspondence should be addressed:** Nicola Grissom, Department of Psychology, University of Minnesota, Minneapolis, MN, 75 East River Road, Minneapolis, MN 55454.

**Keywords:** Sex differences, cognitive flexibility, Set Shift, mice, executive function, touchscreen, rule shifting

## Abstract

Cognitive flexibility can be defined as the ability to adaptively shift between choices or strategies based on environmental feedback and it is disrupted in numerous neuropsychiatric conditions. Individual differences in the computations supporting cognitive flexibility are poised to reveal mechanisms of neuropsychiatric risk and resilience. One critical variable well known to influence individual differences in neuropsychiatric risk is sex. While previous research has identified sex differences in value based decision making in mice, whether sex reflects a major source of variation in cognitive flexibility remains unknown. To directly assess sex-biased individual differences in cognitive flexibility, we developed a novel touchscreen Set Shift task that permits robust and continuous testing in mice. Using this task, we discovered that female mice completed significantly more rule shifts with fewer errors than males. We next employed a suite of computational models that revealed sex-biased individual differences in the computations underlying cognitive flexibility. Overall, our results suggest that following rule shifts, female mice learn the new rule faster and commit to exploiting rule choices sooner compared to males - sometimes because they commit to multiple rules simultaneously. This suggests that increased choice stability in female rodents enhances commitment to a strategy during periods of uncertainty and directly contributes to increased rule shifting. This supports the counterintuitive conclusion that a high degree of stable choice is a strong requirement for enhanced cognitive flexibility in the Set Shift task, one of the gold standard cognitive flexibility tasks.

## Introduction

Cognitive flexibility is a key executive function, and can be defined as the ability to adaptively shift between choices or strategies based on environmental feedback. Maladaptive cognitive flexibility has been implicated in numerous neuropsychiatric conditions, including insufficient flexibility in depression and obsessive compulsive disorder, versus excessive flexibility in autism and psychosis (Addicott et al., 2017; Burguière et al., 2015; Cuthbert and Insel, 2013; Durstewitz et al., 2021; Reimer et al., 2024; Uddin, 2021). Individual differences in the computations supporting cognitive flexibility are thus likely to reveal mechanisms of neuropsychiatric risk and resilience. The gold standard tasks used to probe cognitive flexibility (such as Attentional Set Shifting or the Wisconsin Cart Sorting Test) require both stability during periods of relative certainty and the ability to engage flexible decision making rapidly when the task environment changes (Basu et al., 2023; Bissonette and Powell, 2012; Darrah et al., 2008;

Reimer et al., 2024; Widge et al., 2019). The need to balance stability with flexibility in value-based decision making is better known as the explore-exploit tradeoff, suggesting that individual differences in explore-exploit balance may drive altered cognitive flexibility.

One critical variable well known to influence individual differences in neuropsychiatric risk is sex, including for numerous disorders implicating cognitive flexibility (Grissom et al., 2024; Grissom and Reyes, 2019; Merikangas and Almasy, 2020). Sex differences are highly likely in cognitive flexibility. We have previously found that during value-based decision making tasks, female mice are more likely to both repeatedly execute rewarding choices, and shift decision making strategies sooner following non-rewarded outcomes compared to males (Chen et al., 2021a, 2021b). These findings demonstrate that female mice learn faster during exploration, transitioning to exploit states sooner than males, who are overall more exploratory (Chen et al., 2021b). This is consistent with a larger literature demonstrating stronger outcome encoding in female mice, and conversely reduced sensitivity to risky choices or negative outcomes in male animals (Chen et al., 2021a, 2021b; Cox et al., 2023; Grissom et al., 2024; Grissom and Reyes, 2019; Orsini and Setlow, 2017). These data suggest that sex differences in reward-guided decision making computations, including explore-exploit tradeoffs, may drive distinct approaches to cognitive flexibility tasks.

Here, we demonstrate a newly-developed operant touchscreen Set Shift task that permits robust and continuous testing in mice, and use this task to interrogate sex differences in the computations supporting cognitive flexibility. As in published rat versions of this task (Darrah et al., 2008; Reimer et al., 2024), the mouse Set Shift task employs two different rules and allows repeated, automated, performance based rule shifts without requiring investigator intervention, permitting unlimited rule transitions (up to 200 trials) to allow stronger computational modeling of behavioral strategies. Using this task, we discovered that female mice completed significantly more rule shifts with fewer errors than males. We next employed a suite of computational models to determine what factors drove increased Set Shift performance in females. Consistent with prior literature, we determined via a combined reinforcement learning-drift diffusion model (Reimer et al., 2024) that both choice pre-commitment (bias) and value updating processes (learning rate) are enhanced in female mice on average compared to males. To measure the explore-exploit tradeoff, we used an input-output hidden Markov model, and determined that females more rapidly exit exploratory decision states than males. Critically, female mice were more likely to sometimes exploit multiple rules simultaneously, defined as “ambiguous exploit states” by the hidden Markov model, and this choice strategy was associated with the highest success rates. Collectively, these results suggest enhanced exploit behavior, shown as commitment to (one or more) rules, promotes enhanced performance of cognitive flexibility in female mice.

## Methods

### Animals

Thirty-two BL6129SF1/J mice (16 males and 16 females) were obtained from Jackson Laboratories (stock #101043). Animals arrived at 7 weeks of age and were housed in groups of four. Animals were tattooed for identification and handled before baseline body weights were collected. Two weeks prior to operant chamber training, animals were moved to 0900-2100 h reversed light cycle housing to permit operant testing during the dark period (09:00am-5:00pm). One week prior to operant chamber training, animals were food restricted to 85%-95% of free feeding bodyweight, with *ad libitum* access to water. Animals were pre-exposed to the reinforcer by providing a bottle containing 50% vanilla Ensure for 24 h in the home cage, verifying consumption by all cagemates. Starting at 12 weeks of age, behavioral training and testing in the operant chambers occurred five days per week (Monday-Friday), and animals were fed each day following behavior, with *ad libitum* food access provided on Fridays. Behavioral training and testing occurred in the same chamber for each animal and operant chambers were located in unique female and male behavior rooms to avoid potential confounds. One female animal unexpectedly died before Set Shift training began due to natural causes, producing a final n of 31 mice (16 males and 15 females). All animals were cared for in accordance with the National Institution of Health and the University of Minnesota Institutional Animal Care and Use Committee guidelines.

### Apparatus

Sixteen triangular touchscreen operant chambers (Bussey-Saksida design, Lafayette Instrument Co., Lafayette, IN), enclosed inside sound attenuating cabinets were used for behavioral training and testing. Two of the three chamber walls were black, acrylic plastic. The third wall housed the touchscreen and was positioned directly opposite of the reward port and magazine. A two-hole mask over the touchscreen defined two possible choice options. The magazine provided Ensure liquid reinforcer, delivered by a peristaltic pump (280 ms pump duration, ∼7ul). ABET-Cognition software (Lafayette Instrument Co., Lafayette, IN) was used to program operant behavioral schedules and to collect all training and testing data.

### Pre-training

After habituation to the operant chamber, animals were exposed to daily touchscreen training schedules including Initial Touch, Must Touch, and Must Initiate as previously described (Chen et al., 2021a). As part of the touchscreen operant training routine, animals also completed the Punish Incorrect schedule but the stimuli displayed on the touchscreen during the task was adapted to be consistent with stimuli presented during the Set Shift task. After mice reached criterion on each schedule (two consecutive days with 30 trials completed in 30 minutes or two consecutive days with 60 trials completed in 60 minutes, depending on schedule), they moved on to the next schedule. On average, animals spent five days on each schedule.

### Rule Shaping

To control for training order effects, animals were assigned to learn either the “light” rule (n=8, 4F, 4M) or the “side” rule (left or right) (n=7, 3F, 4M) first, ensuring that cagemates of each sex were evenly distributed across conditions (light-left-right; light-right-left; left-right-light; right-left-light) (**Supp. Figure 1a**). Rule shaping schedules were rewarded deterministically. During Light Shaping, animals only received reward for choosing the “light” image (illuminated square), while during Left Shaping and Right Shaping, animals were only rewarded for choosing the left option or the right option, respectively. Animals spent five days on each schedule, and each animal reached the shaping performance criterion of completing at least 10 consecutive correct trials, confirming that all animals learned each individual rule.

### Touchscreen Set Shift Task for mice

The touchscreen operant Set Shift task (**Figure 1a**) was modified from standard operant chamber designs in de Oliveira et al., 2021 and Reimer, et al. 2024 and originally developed by Darrah et al., 2008 (Darrah et al., 2008; de Oliveira et al., 2021; Reimer et al., 2024). To probe putative cognitive flexibility, the Set Shift task employs two different rules–a light rule where the animal must choose the “light” (illuminated square) regardless of what side it appears on, and a side rule where the animal must choose a side (left or right) regardless of the light location (**Figure 1a**). If the animal does not make a choice within the response time limit (normally 10 seconds, but 3 second response time limits were also tested as described in Results), an omission is logged. Following five consecutive correct choices without omission in a given rule, the criterion is met and the rewarded rule shifts to a different rule without warning. The lack of reward for a previous rule response is the only indication that the rewarded rule has shifted.

**Figure 1.**
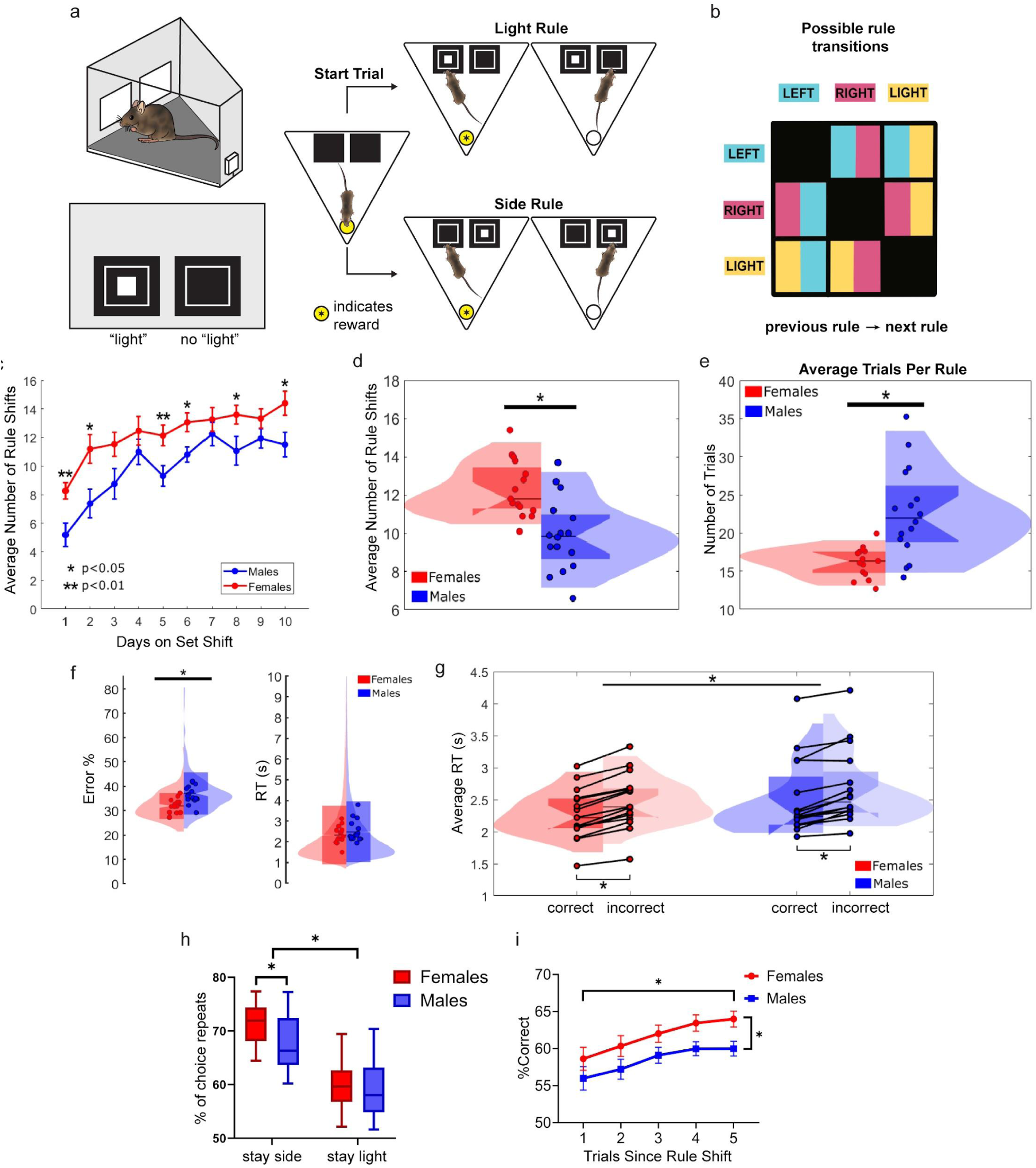
(a) The mouse operant touchscreen Set Shift task. In the images displayed on the touchscreen, the outer white square indicates that the touch/choice is active while the inner white illuminated square is the “light” cute. (b) All possible rule transitions. Rules are selected in a random equal-number way. (c) Though Set Shift performance (number of completed rule shifts) improves for all animals across the 10 total days on Set Shift (p=8.7981e-12), female mice consistently complete more average rule shifts compared to males across daily sessions (p=9.4899e-05) (n=31, 15F/16M, ∼200 trials/session/mouse). (d) Averaging across Set Shift days, female mice complete significantly more rule shifts than males (p=0.0005) (n=31, 15F/16M, ∼2000 trials/mouse). (e) Female mice reach criterion (5 consecutive correct choices without omissions) in fewer trials than males on average, spending less time in each rule and rule shifting faster than males (p=0.0004). (f) Female mice demonstrate fewer errors than males during Set Shift (p=0.00075) but there are no significant sex differences in overall, average response time (RT). (g) Considering how RT differs by performance across sex reveals a significant main effect of performance (correct/incorrect) (p=8.0876e-286) and a significant interaction effect between sex and performance (p=0.0005). (h) When the previous choice (side or light) was correct, animals more frequently repeated the previously correct side choice than the previously correct light choice (p<0.05). On average, female mice repeated the previously correct side choice even more frequently than males (p = 0.0351), demonstrating enhanced correct choice repetition. (i) All animals get more choices correct as trials increase following a rule shift (p = 0.0006), but female mice make significantly more correct choices across trials following rule shifts compared to males (p<0.0001).

During Set Shift, animals are required to inhibit a prepotent response by exploiting the current rule and exploring to learn the new rule after the environment “shifts.” Unique to this Set Shift task, the rewarded rule shifts to any of the other rules (six total transition combinations (**Figure 1b**), allowing for inter- and extradimensional shifts. The next rule is selected in a random-equal-number way to prevent predictability. Each trial is initiated at the magazine when the tray light turns on. Following initiation, animals have a response time limit (10-3 s depending on schedule) during which they must execute a choice before a three-second timeout begins. Incorrect choices similarly evoke this timeout, paired with an incorrect tone (3000 Hz). After each choice or omission, a three-second inter-trial interval (ITI) exists before animals can initiate the next trial. Touches during the ITI are also recorded. Immediately following a correct choice, the feeder pump distributes liquid reward (Ensure) and simultaneously the tray light turns on. Once reward is collected, the tray light turns off and the ITI starts. The Set Shift task lasts for 200 trials or 60 minutes, whichever occurs first. Throughout the task, mice are allowed to complete as many rule shifts as they can in the allotted time.

### Set Shift schedule modifications to test reduced response time limits in mice (3 seconds)

As a new task and with the desire to detect sex-biased individual differences in behavior, the baseline Set Shift task was as described above, but allowed a 10 second response time limit. To assess the ability of mice to perform the task with the same response time limit as rats in a standard operant chamber (Darrah et al., 2008; Reimer et al., 2024), a second Set Shift schedule was created with a response time limit of three seconds. An additional descending response time limit schedule was developed to further investigate how the response time limit influenced female and male mouse behavior during Set Shift. This schedule started with a 10 second response time limit that decreased by one second after each completed rule shift until the response time limit reached three seconds where it remained for the duration of the session (see Results). Given that the response time limit decreased with each completed rule shift, not all animals reached the three second response time limit in each session. One female and two males only reached the three second response time limit during some sessions of this schedule. In those cases, the number of rule shifts completed during the three second response time limit was recorded as zero. One male animal never reached the three second response time limit during this schedule and as such, could not contribute data to the analysis of this schedule.

### Data Analysis

Primary behavioral analyses were conducted using MATLAB version 2022b, Python version 3.11, and GraphPad Prism 10. Generalized linear mixed effects models (GLMMs) (see below) and repeated measures analysis of covariance were used for statistical testing. Animal counts were predetermined by a power analysis to detect the standard effect size of sex differences common in previous publications from the lab (Chen et al., 2021a, 2021b), targeting 80% power (G*Power 3.1).

Consistent with previous studies (Reimer et al., 2024), repeated-measures data including response times, accuracy, and number of completed rule shifts were analyzed with GLMMs as gamma regressions for response times, and linear regressions for completed rule shifts and errors/performance. The general model for these analyses was:

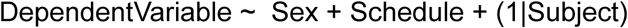

Sex differences in the average number of trials to rule shift was statistically compared using a repeated-measures ANOVA.

### Reinforcement Learning Drift Diffusion Model (RLDDM)

We used a reinforcement learning drift diffusion model (RLDDM) to model decision making in the Set Shift task (Reimer et al., 2024). This model combines elements of two prominent computational modeling approaches for decision making: 1) a reinforcement learning model, which updates trial-to-trial value learning, and 2) a drift diffusion model, which models evidence accumulation up to choice. The RLDDM was fit to mouse Set Shift behavior (the sequences of choices and response times each session) using Markov-Chain Monte Carlo with the HDDM Python package (Reimer et al., 2024; Wiecki et al., 2013). To ensure reliable estimation of parameters, four independent chains (2,000 total samples) were run for each model and convergence was confirmed by assessing whether the Gelman-Rubin statistic was <1.1 (Gelman and Rubin, 1992; Reimer et al., 2024). To examine sex-based influence, we allowed the model to fit different intercept terms for males and females, while assuming similar variability across sexes. This allows us to evaluate how strongly each model parameter (boundary separation, drift rate, bias, non-decision time, learning rate, forgetfulness, and surprise) is affected by sex. Using the RLDDM, values are assigned to each choice based on estimates of the current side (left and right) and “light” values. The model uses those values to generate a choice using a drift diffusion process, accumulating evidence until the decision boundary is reached. As such, trials with a larger total difference in value between choices are more likely to have shorter response times and more consistent choices. Depending on whether a choice is rewarded, values are updated using a reinforcement learning process. Modeling the data from each group (females and males), the posterior distribution of group model parameter differences inform which computations show the largest sex biases. The full set of equations for the RLDDM is as follows:

The value of a choice (V) is the sum of its respective side and light values (Q):

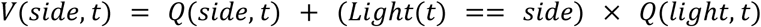

Drift (*v*(*t*)) on each trial is directed proportionally to the side with the highest value:

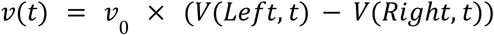

Bias (β(*t*)) on each trial is directed proportionally to the side with the highest value, ignoring the light:

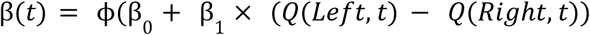

On each trial, non-decision time (τ(*t*)) was sampled from a uniform distribution with mean τ_0_ and width τ_*var*_:

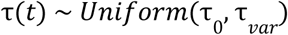

For each trial, response times (RT) follow a Wiener First Passage Time (WFPT) distribution with parameters for drift (*v*(*t*)), boundary separation (α), bias (β(*t*)), and non-decision time (τ(*t*)):

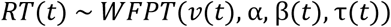

Separate non-linear learning rates (δ) were calculated for each chosen feature (*C*) to model surprise (γ):

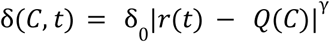

The value of each chosen feature was updated via a standard Rescorla-Wagner learning rule:

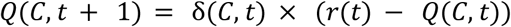

The value of unchosen features diminishes according to a forgetfulness factor (*F*):

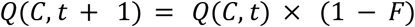

The RLDDM has 10 parameters in total: initial values for the side and light features, drift rate (*v*_0_), baseline bias (β_0_), non-decision time (τ_0_), variability of non-decision time (τ_*var*_), boundary separation (α), baseline learning rate (δ_0_), surprise (γ), and forgetfulness (*F*). Parameters were estimated using a Bayesian hierarchical approach. To determine the effect of sex on model parameters, the group-level distributions for the parameters were assessed using the probability of direction (PD) and the region of practical equivalence (ROPE) (Makowski et al., 2019; Reimer et al., 2024). PD refers to the proportion of the parameter distribution greater than 0 with a value above 0.5 (indicating a general increase in the parameter across groups) and a value below o.5 (indicating a general decrease in the parameter across groups). ROPE refers to the proportion of the parameter distribution that falls within a region that is practically equivalent to a null-effect (+/- 0.1). PD indicates the existence and direction of an effect while ROPE establishes significance.

### Input-Output hidden Markov model (ioHMM)

An input-output hidden Markov model (ioHMM) was used to label latent cognitive states underlying Set Shift behavior. The HMM framework assumes that choices are generated from some unobserved latent cognitive states. Our model identified two general types of cognitive states–exploration and exploitation–defined by their unique patterns of choice. Here, animal’s choice behaviors are modeled as emissions (observations) from one of the four distinct latent cognitive states - explore, exploit left, exploit right, and exploit light. In the explore state, the emission probability for specific choice types are uniform, meaning that during the explore state, the generation of all choice types (left, right, and light choice) are equally probable. The exploit states only emit the type of choice being exploited (e.g. the exploit left state only generates left choices). The transition matrix fit to the subject captures the unique transition probability between these four states in each individual. To disambiguate between choice dimensions (side and light), the location of the light cue was used as an input layer into the ioHMM. Together, the final ioHMM receives each animals’ choice sequences and the respective locations of the light cue, and outputs the most probable latent state for each trial.

The base ioHMM MATLAB code was adapted from Ebitz et al., 2019 and Ebitz et al., 2020 (Ebitz et al., 2020, 2019), and Kevin Murphy’s MATLAB HMM toolbox. Below is a detailed description of the ioHMM:

Within the HMM framework, choices or “emissions” (y) are generated by an unobserved decision process within some hidden, latent state (z). These latent states are defined by both the probability of making each choice (k) out of (N_k_) possible options given the state of the system (e.g. whether the light is on the left or right side) and the probability of transitioning from each state to every other state. This framework was extended to allow inputs, such as the location of the light, to influence the probability of observing each emission in each state (Bengio and Frasconi, 1994). These emission probabilities (**Figure 3a**) differ across the two broad types of states in this model–the explore state and the rule states–based on the differences in their behavioral dynamics: one representing more random choices made during exploration and the other involving consistent, repeated choices following a given rule. For example, during exploration, animals have an equal, but random probability of making left and right choices regardless of the light location.

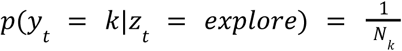

However, the location of the light is critical for disambiguating between the different rule states. For example, if an animal is following the left rule, they will repeatedly make left choices regardless of whether the light is on the left or right side. However, if an animal is exploiting the light rule, they will only make left choices when the light is on the left side and right choices when the light is on the right side. That is, the observation model for rule states was dependent on whether or not the observation (choice + light location) met the condition for the current rule:

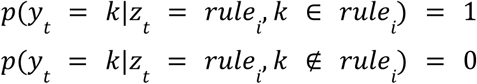

The latent states in this model are Markovian, meaning that they depend only on the most recent state (z_t_) and the most recent location of the light (l_t_), and are independent of time:

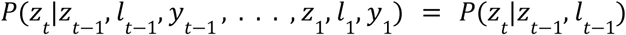

The probabilities of each state transition were described by the one-time-step probability of transitioning between every combination of past and future states (i,j). Unlike the emissions matrix, the transition matrix was not influenced by the location of the light.

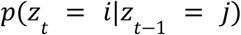

The model had four possible states (one explore state and three rule (exploit) states). Parameters were tied across exploit states such that each exploit state had the same probability of beginning from exploration and of sustaining itself. Transitions from exploration to the exploitative states were similarly tied, making it equally likely to start exploiting any of the three rule states after exploration. Rule sequences were pseudo-randomized to prevent prediction of the next rule, including the starting rule. As such, transitions directly between exploit states were not permitted and choices were initialized in the explore state. The model assumed that mice had to pass through the explore state in order to gain enough information to start exploiting a new rule, even if only for a single trial. This modeling choice also reduced the number of free parameters to estimate, allowing us to estimate fewer parameters more accurately with a limited number of trials. As such, the model had two free parameters.

The model was fit via expectation-maximization using the Baum Welch algorithm (Blimes, 1998). This algorithm identifies a (possibly local) maxima of the complete-data likelihood based on the joint probability of the latent state sequence and the sequence of observed choices. The algorithm was reinitialized with random seeds 20 times, and the model that maximized the observed (incomplete) data log likelihood was selected as the best for each session. To label latent states from choices, the Viterbi algorithm was used to identify the most probable *a posteriori* sequence of latent states (Murphy, 2021; Rabiner, 1989).

## Results

### Female mice show enhanced Set Shifting compared to males

Age-matched female and male wildtype mice (BL6129SF1/J, n=31, 15F, 16M) were trained to perform a novel Set Shift task designed for mouse touchscreen operant chambers. To uncover computational mechanisms for sex differences in cognitive flexibility, we adapted a gold-standard rat operant set shifting task for a mouse touchscreen (**Figure 1a**). This type of Set Shift task employs two different rules–a light rule where the animal must choose the “light” (illuminated inner square) regardless of what side it appears on, and a side rule where the animal must choose a side (left or right) regardless of the “light” location. Following five consecutive correct choices without omissions, the rule changes with no cues or indications that it has done so (Reimer et al., 2024). To better enable computational modeling of individual differences in behavioral performance in mice, we made three key modifications from the standard task. The standard task has 1) a fixed *sequence* of rules, with extra-dimensional rule shifts only (always shifting between a light and a side rule), and 2) a fixed *number* of rules and transitions (Darrah et al., 2008; Reimer et al., 2024). Here, we 1) allowed rule *sequences* to vary in a random-equal-number way, permitting both inter- and extra-dimensional rule shifts (six total transition combinations (**Figure 1b**)), and 2) did not limit the number of rule shifts completed during daily sessions (200 trials or 60 minutes). We also 3) extended the decision time to 10 seconds consistent with other mouse touchscreen continuous performance tasks (Grissom et al., 2015; Horner et al., 2013; Mar et al., 2013). This design allowed us to measure sex differences in Set Shift performance as well as the contributions of different task features in driving these differences.

Prior to testing on the Set Shift task, animals were trained on each rule separately in a counterbalanced order. To control for bias produced by the order in which animals were exposed to each rule, cage-mates of each sex were evenly distributed to each training order–with animals learning either the Light Rule, Right Rule, or Left rule first (**Supp. Figure 1a**). All animals successfully learned to perform the Set Shift task and completed within-session rule shifts, regardless of training order (**Supp. Figure 1b**).

Following exposure to each rule individually, testing began on the full Set Shift task including all three rules. A robust sex-biased individual difference in the average number of rule shifts completed emerged on the first day of the full Set Shift task (**Figure 1c**). Training order also influenced the number of rule shifts completed on the first day of Set Shift. Animals that completed side shaping last (immediately preceding the full Set Shift task) performed significantly fewer rule shifts compared to animals that completed light shaping last (**Supp. Figure 2a-b**). This early training order effect also interacted with sex differences. Male mice that completed side shaping last performed the fewest number of rule shifts on the first day of Set Shift (**Supp. Figure 2c**) (Shaping rule effect: p=0.0022, sex effect: p=0.0004). Despite the initial presence of these training order effects, they did not persist past the first day on the full Set Shift task. Though all animals’ Set Shift performance improved across the 10 days on the task, female mice consistently completed more rule shifts on average compared to males (**Figure 1c**) (n=31, 15F/16M, ∼200 trials/session, 10 sessions/animal, GLMM main effect of session: p=8.7981e-12 & main effect of sex: p=9.4899e-05).

To examine effects driving this robust main effect, we collated Set Shift performance across 10 days. Averaging across sessions reveals robust sex-biased individual differences in rule shifting performance where female mice complete more rule shifts (n=31, 15F/16M, ∼2000 trials/animal, p=0.0005), meeting criterion to shift in fewer trials than males (p=0.0004) (**Figure 1d-e**).

Multiple factors contribute to the number of rule shifts animals can perform. Female mice may complete more rule shifts than males because they are more accurate during the task, or because female mice are simply faster at the task. To test these two hypotheses, we assessed average error percentages and response times (RT) across sessions and identified that female mice make fewer errors than males during Set Shift (p<0.00075), with no significant differences in overall average RT (**Figure 1f**). Though RT does not differ across sex alone, average RT is sensitive to performance (correct and incorrect trials). Splitting RT by sex and performance revealed that RTs during correct trials were significantly faster than RTs during incorrect trials at both the individual and group levels (GLMM, gamma regression, p=8.0876e-286) however, this effect was stronger in the female mice (GLMM, gamma regression, sex*performance interaction, p=0.0005)(**Figure 1g**). This suggests that the sex-biased individual differences present in the number of completed rule shifts emerge not simply due to females completing trials faster than males, but due to differences in trial accuracy.

Previous research has identified sex differences in decision making strategy (Chen et al., 2021a, 2021b) and outcome sensitivity (Chen et al., 2021a, 2021b; Cox et al., 2023; Grissom et al., 2024; Grissom and Reyes, 2019; Orsini and Setlow, 2017) in rodents. Consistent with these concepts, not only did female mice make more correct choices than males during the Set Shift task, they used trial feedback (whether or not they were rewarded) differently. Though all animals more frequently repeated the previously correct side choice than the previously correct light choice (p=1.34012e-11), females demonstrated significantly higher correct choice repetition compared to males (p = 0.0351) (**Figure 1h**). Further, when aligned to the trials (1-5) following rule shifts, though percent correct increases across the trials following a rule shift in all animals (p = 0.0006), female mice are more accurate across trials immediately following rule shifts (p=6.47705e-05) (**Figure 1i**). Overall, higher accuracy and increased correct choice repetition in female mice both contributed to females completing more rule shifts than males. These findings suggest that female mice may differ from males in the computations used in choice selection and reward learning.

### Stronger effects of choice bias and learning rate in female mice

Several broad computational mechanisms might simultaneously contribute to the sex differences we observe in cognitive flexibility. Differential reward updating (Chen et al., 2021b; Golden et al., 2023; Palmer et al., 2024), action repetition (Chen et al., 2021a), and outcome sensitivity (Cox et al., 2023; Palmer et al., 2024) across sex have all been shown to influence sex differences in decision making. Our analyses thus far suggest that female mice repeat correct choices more frequently, pointing to contributions of value updating and/or action repetition as a key variable driving sex-biased Set Shift behavior. To get a global evaluation of the multiple potential mechanisms contributing to sex differences in cognitive flexibility, we turned to an innovative combination reinforcement learning drift diffusion model (RLDDM) (**Figure 2a**). With both reinforcement learning and perceptual decision making components, the RLDDM is well suited to model Set Shift behavior (Reimer et al., 2024).

**Figure 2.**
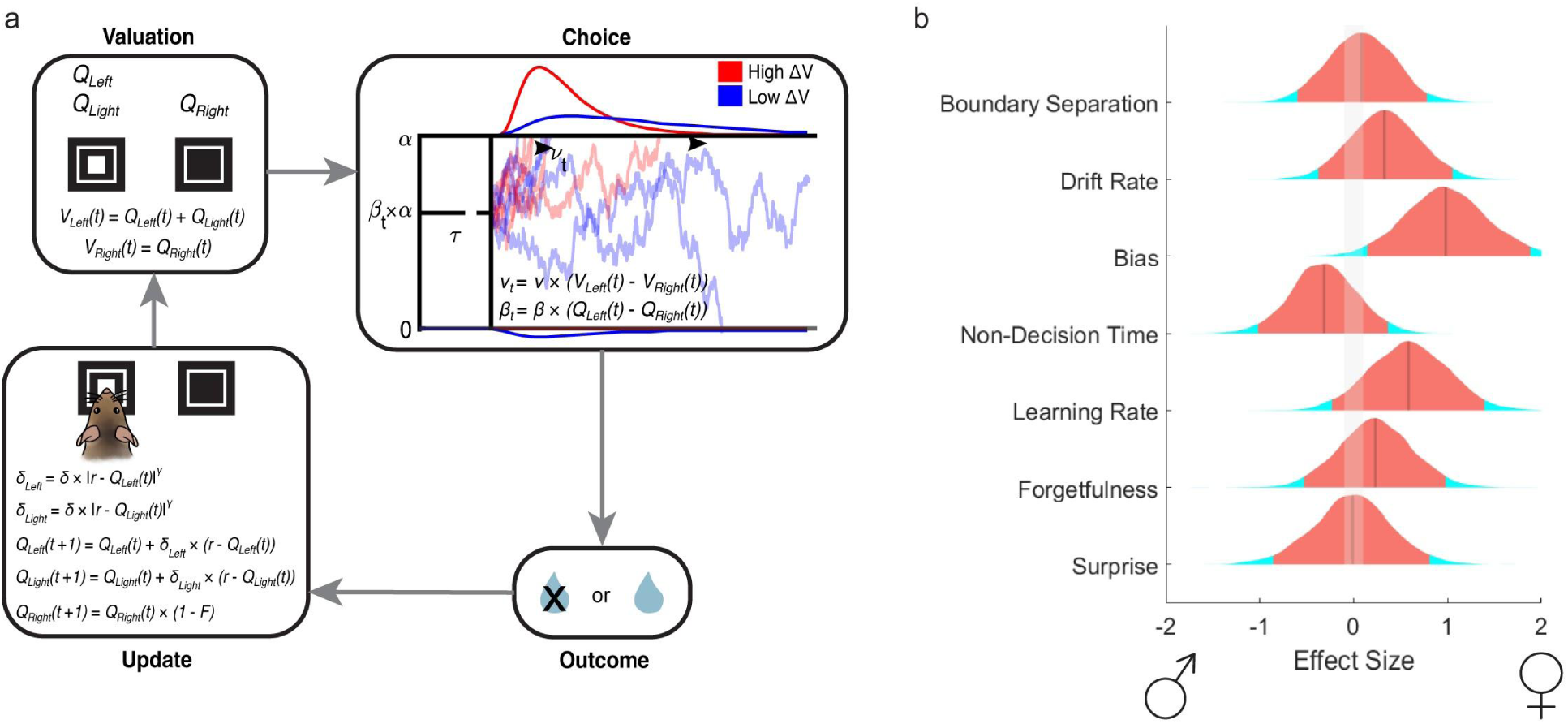
(a) Schematic of the reinforcement learning drift diffusion model (RLDDM). (b) Distributions of the effect of sex on each model parameter across 4000 posterior draws. The 95% highest density interval is depicted in red/orange, the solid gray line indicates the median of the distribution, and the shaded gray area on either side of 0 represents the region of practical equivalence (ROPE) for a null effect (effect size < 0.1). Effect size by sex is indicated by the x-axis, with 2 representing a stronger effect in females and −2 representing a stronger effect in males.

The RLDDM (**Figure 2a**) was fit to female and male Set Shift behavior, allowing us to evaluate how strongly each model parameter (boundary separation, drift rate, bias, non-decision time, learning rate, forgetfulness, and surprise) is affected by sex. Boundary separation reflects the amount of evidence required to make a choice. Drift rate captures the efficacy of evidence accumulation–how quickly the model is driven toward a correct choice. The bias term represents the ability to pre-commit to the higher value choice based on the learned value differences between choice options. Non-decision time captures the speed of sensorimotor processing–the time it takes to process the stimulus and execute a motor (choice) response. Learning rate (value updating) and forgetfulness (decay of the unchosen option) were included as part of the reinforcement learning component of the model. Overall, the RLDDM accurately modeled mouse Set Shift behavior (**Supp. Figure 3**).

Corroborating prior results, two main parameters associated with value updating and choice repetition were the strongest contributors to the sex differences observed in Set Shift behavior. Effects of learning rate and bias (pre-commitment to the higher value choice) were stronger in female mice compared to males (**Figure 2b**). This bias effect indicates that once the current rule choice is learned, female mice more strongly commit to that learned rule response. Together, these data suggest that computationally, female mice demonstrate increased cognitive flexibility during Set Shift by a combination of increased updating from learned values and repetition of higher value (rule) choices.

### Faster transitions out of exploration following rule shifts in females

Our behavioral analyses and RLDDM findings broadly indicate that female mice repeatedly choose high value options throughout the Set Shift task. How are they achieving this, given that the correct response is continually shifting? This task requires animals to both commit to a rule-specific choice during periods of relative certainty (stable rule responding) and to flexibly explore alternative options during periods of relative uncertainty (rule shifts). This suggests that animals will occupy different cognitive states as they navigate the unsignaled changes in this environment–exploiting the current rule state and exploring to learn the new rule state following a rule shift (Ebitz et al., 2020, 2019). Cognitive flexibility can be formulated as the rate of adaptive transitions between exploration and exploitation. The most optimal strategy for the task is to rapidly transition to exploration following a loss on a previous rule, and transition to exploiting a new rule as rapidly as possible. Therefore, we hypothesized that female mice were faster at one or both of these processes. We have previously identified sex differences in explore-exploit state occupancy in a restless bandit task, where female mice exit exploration sooner than males because they learn about rewarding choices faster (Chen et al., 2021a, 2021b). Our RLDDM findings similarly indicate a higher learning rate in females during Set Shift, further informing our hypothesis. Our prior data suggests that there should be no difference in the ability to become flexible (exploit→explore), but the ability to transition away from exploratory flexibility into a new rule state (explore→exploit) should be enhanced in females.

To identify latent cognitive states underlying Set Shift behavior, an input-output hidden Markov model (ioHMM) structure (**Figure 3a**) was used, providing both behavioral information about the choices animals were making (left or right), as well as task information about the location of the light (left or right) to the model. With this input, the model estimates the most probable states that an animal is in on each trial. Our ioHMM models two types of cognitive states – exploration across rules, and exploitation of a certain rule (left, right, or light).

**Figure 3.**
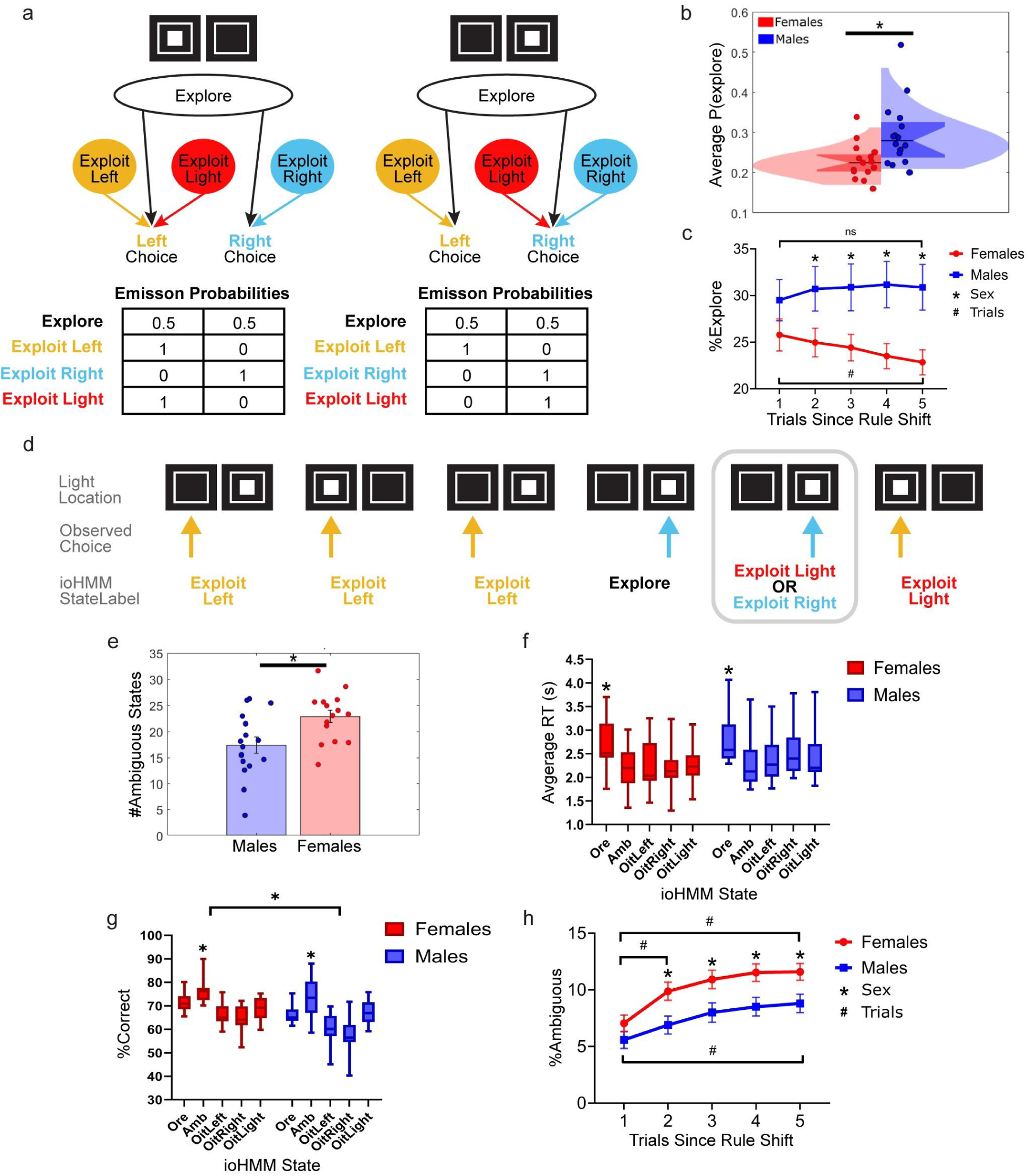
(a) Schematic of the input-output hidden Markov model (ioHMM), illustrating how the emission probabilities for making a left or right choice in each state change in response to the input (location of the light stimulus). (b) On average, male mice have a higher probability of being in the explore state compared to females throughout the Set Shift task (p = 0.0083). (c) One to five trials after rule shifts, the average percent of explore labeled trials significantly decreases in female mice (p=0.0432), while male mice remain in the explore state across trials following rule shifts, presenting a sex difference in the percent of explore labeled trials for trials 2-5 after rule shifts (p=2.19611e-06). (d) Example schematic of an ambiguous state (outlined in light grey) occurring at trial t following an explore labeled state (t-1), where the animal’s choice and the location of the light remained unchanged from trials t-1 to trial t. The ioHMM gave an equal probability of two exploit states of different dimensions–at trial t, the animal could have been in the exploit light or exploit right state. At trial t+1, the light stimulus moves and the animal chooses the side where the light is present, resulting in the ioHMM labeling trial t+1 as an exploit light trial. (e) On average, female mice have significantly more ambiguous state trials (trials with two equiprobable states (an exploit side state and an exploit light state) compared to males (p=0.01). (f) For female and male mice, response times (RT) were significantly slower during explore labeled states (p=0.0009), compared to exploit labeled states and ambiguous states, which were similar in RT to exploit states. (g) For female and male mice, accuracy (average percent correct) was higher for ambiguous state trials compared to explore and exploit labeled trials (p=8.11006e-17). Overall, percent correct was higher in female mice regardless of state occupancy (p=6.46393e-07). (h) In all animals, the average percent of ambiguous trials significantly increased across trials following rule shifts (p=5.77367E-06). In female mice only, the percent of ambiguous trials increased from 1-2 trials following a rule shift (p=0.0138) and were overall higher on average compared to males (p=4.85193e-07).

Male mice overall spent more time in exploration during Set Shift compared to females (**Figure 3b**). To test the hypothesis that female mice specifically show increased transitions from exploration to exploitation, we quantified the percent of explore-labeled trials for the first five trials following rule shifts. Female mice, but not males, significantly decreased their exploration from the first to the fifth trial (p=0.0432), while males did not change their exploration over these trials and were significantly different from females on trials 2 through 5 (p=2.19611e-06) (**Figure 3c**). Given that the first erroneous choice in a new rule is the only indication the mice receive that the rule has changed, we would not expect internal states to change until after the first trial in a new rule (at the earliest). Consistent with this, there was no significant difference in exploration between females and males during the first trial after a rule shift. However, females demonstrated a significantly lower average percentage of explore labeled trials than males trials 2-5 following rule shifts as their respective strategies further diverged across trials (**Figure 3c**). This suggests that the ability to transition away from exploration in fewer trials largely contributes to the increased number of completed rule shifts observed in female mice.

### Simultaneous exploitation of multiple rule states in females

In the Set Shift task, transitions away from exploration imply transitions towards exploitation of specific rules. However, it would take multiple trials to test all possible rule states, because each trial contains two response options, as well as two dimensions (side and light) to disambiguate. The rapid transitions into exploitation for females may not allow for adequate trials to fully disambiguate the rule. One possibility is that females may exploit multiple rules simultaneously.

With the external rule identity hidden from both the animal and the ioHMM, our exploit states are not mutually exclusive, allowing us to observe whether or not they are co-occupied (occurring in equal probability). On an overall average of 10.39 percent of trials, the ioHMM gave an equal probability of the animal being in two states. These “ambiguous states” always occurred between the exploit light state and an exploit side state (**Figure 3d**).

Consistent with the idea that female mice may exploit multiple rules simultaneously, ambiguous states occurred more frequently for female mice (avg. 22.85 trials/session) than male mice overall (avg. 17.35 trials/session) (p=0.01) (**Figure 3e**). While ambiguous states may represent an ‘in-between’ state where an animal is not exploiting a single rule, they are not explore states, as estimated by the model. This is further supported by assessing average RT and accuracy in each state. Given the relationship between cognitive states, decision making confidence, and choice accuracy, previous research has identified that exploratory choices are slower and less accurate than exploitative choices (Chen et al., 2021b). Consistent with previous findings, RTs in explore-labeled trials are significantly slower than RTs in exploit states, regardless of sex (p=0.0009) (**Figure 3f**). Further, RTs in ambiguous state trials did not significantly differ from RTs in other exploit states, providing evidence that ambiguous states are a type of exploit state. As an exploit state, ambiguous trials showed significantly higher accuracy compared to explore trials (p=8.11006e-17) (**Figure 3g**). Additionally, accuracy was higher during ambiguous state trials compared to trials in all types of exploit states (left, right, and light). This argues that ambiguous exploit state trials are unique, but the increased success of ambiguous state trials is not necessarily surprising (see Discussion).

Thus far, our analysis of ambiguous states suggests that these states do indeed represent trials where animals co-occupy two unique exploit states. To assess whether female mice transition to exploitation in fewer trials than males following rule shifts by simultaneously exploiting two states, we quantified the percent of ambiguous labeled states 1-5 trials following rule shifts–similar to our analysis of exploration following rule shifts. While the percent of explore labeled trials decreases in females 1-5 trials following rule shifts, the percent of ambiguous state trials increases 1-5 trials following rule shifts in all animals (p=5.77367E-06), with an overall higher occurrence in females (p=4.85193e-07) (**Figure 3h**). This suggests that co-occupancy of two exploit states exists as a strategy to navigate periods of relative uncertainty during the Set Shift task and importantly, that female mice use this strategy more frequently than males.

In an effort to validate our ioHMM results, we analyzed and compared shuffled trial-by-trial state data (**Supp. Figure 4**). Shuffling the order that states occurred in eliminated trial specific effects across measures. This method unpaired state labels from their assigned trials, eliminating state-specific differences in RT and accuracy (**Supp. Figure 4a-b**). As expected, the shuffling did not change overall sex-biased differences in these measures, but did eliminate temporal effects (**Supp. Figure 4c-d**). This further validates that the state assignments and measures we observe are not random effects, but rather reflect latent cognitive patterns/states.

Overall, our ioHMM findings indicate that sex differences in Set Shift performance are largely influenced by differences in decision making strategy. When deliberating between rules during periods of uncertainty, female mice transition to exploitation more quickly than males by simultaneously exploiting two rule states.

### Task design impacts mouse Set Shift performance and disturbs previously observed sex differences

The initial version of the Set Shift task allowed 10 seconds after stimulus display for the animals to make a choice. If the animal did not make a choice within the 10 second response time limit, the trial would timeout, no reward would be received, and the next trial could be initiated following a three second inter-trial-interval. The Set Shift task for rats typically allows a three second response time limit (Reimer et al., 2024). After collecting 10 days of testing on the 10 second response limit Set Shift task, we also tested two versions of the task where response time limits dropped to three seconds (**Supp. Figure 5**). Decreasing response time limits dramatically reduced successful rule shifts in all mice, and temporarily reduced differences in performance across sexes. Moving animals back to a 10 second response time limit immediately improved performance (**Supp. Figure 5c**), suggesting that shortened deliberation times were deleterious to the cognitive strategies all animals used in the task, although animals were making decisions faster than three seconds in the original task (compare **Figure 1f**). Finally, we implemented a version where response time limits shortened from 10 seconds to three seconds with each successful rule shift. Despite the slower transition to faster response time limits, performance at the three second response time limit was not improved compared to the base three second schedule and the sex difference did not reemerge (**Supp. Figure 5e-g**). These data indicate that having adequate time to deliberate is a critical contributor to the sex differences we see in Set Shift behavior, consistent with our modeling results above.

## Discussion

Individual differences in the computations supporting cognitive flexibility are poised to reveal mechanisms of neuropsychiatric risk and resilience. One critical variable well known to influence neuropsychiatric risk is sex, and sex differences have long been seen in more general decision making tasks. To directly assess sex-biased individual differences in cognitive flexibility, we developed a novel touchscreen Set Shift task for mice during which female mice demonstrate enhanced cognitive flexibility by completing more rule shifts with fewer errors than males. Using computational modeling, we revealed sex-biased individual differences in the computations underlying cognitive flexibility. Overall, our results suggest that following rule shifts, female mice learn the new rule faster and commit to exploiting rule choices sooner compared to males - sometimes because they commit to multiple rules simultaneously. This suggests that increased choice stability in female rodents enhances commitment to a strategy during periods of uncertainty and directly contributes to increased Set Shifting. Our data support the somewhat surprising conclusion that a tendency towards exploiting options in decision making can drive enhanced cognitive flexibility in an operant Set Shift task, one of the gold standard cognitive flexibility tasks.

Our findings corroborate numerous prior findings of sex differences in the computations supporting decision making generally, and point to enhanced choice stability and reduced exploration in females compared to males. Prior work from our lab identified faster learning rates in female mice using a spatial restless bandit task (Chen et al., 2021b), as well as stronger side choice biases in female mice using a visual restless bandit task where animals had to learn image-value associations (Chen et al., 2021a). Here, using a reinforcement learning drift diffusion model (RLDDM) (Reimer et al., 2024), we similarly find stronger effects of learning rate and bias/pre-commitment in female mice. Further, using a hidden Markov model to understand explore-exploit bias in the restless bandit task, our lab has previously identified that female mice learn faster during exploration, transitioning to exploit states sooner than males who are overall more exploratory (Chen et al., 2021b). In the current experiment, we again found increased exploration in male mice, and that female mice were faster to transition into exploitation compared to males. Our findings are also consistent with a broader literature showing sex differences in outcome sensitivity and learning rate during decision making. Compared to males, female mice demonstrate stronger sensitivity to negative reward outcomes and increased strategy switching following non-rewarded trials (Cox et al., 2023). Likewise, sensitivity to risk and negative reward outcomes is decreased in male rodents (Cox et al., 2023; Orsini et al., 2022). These effects may be related to sex differences in the dopamine system that are partially regulated by estrogen signaling (Golden et al., 2023; Yoest et al., 2018) but also show evidence of sex chromosome regulation (Grissom et al., 2024). Indeed, male mice also show greater variability than females in untrained behavior without a role for estrous cycling (Levy et al., 2023). Collectively, multiple lines of evidence suggest that sex differences in multiple neural systems may be responsible for repeated findings that females are more stable and committed in action selection and decision making (Grissom et al., 2024).

Following rule shifts, we found that female mice were faster to transition into exploitation - meaning commitment to a particular response strategy. Intriguingly, these exploit states were also more likely to be “ambiguous”-trials where the model gave an equal probability of the animal exploiting a side state and the light state. In other words, the behavior of the animal was consistent with multiple rule states. A key question is whether these states are artifacts of the model, or reflect actual computational and neural engagement of multiple rules at once, especially in females compared to males. We find that trials marked as ambiguous states are behaviorally distinct from explore states. Ambiguous exploit state response times are as fast as when animals make single-rule exploit choices, and faster than explore trials in both sexes, consistent with the idea that exploration is more cognitively demanding than exploit decisions (Chen et al., 2021b; Ebitz et al., 2019, 2018). Surprisingly, the accuracy of ambiguous exploit state choices are also *higher* than any other state. However, being in an ambiguous state may lead to more success because they allow for commitment to two of the possible rules without needing to know the *specific* rule. By definition, ambiguous states occur during trials where the potential side rule and light rule are congruent, which are “easier” than incongruent trials where the light location is in conflict with the side an animal wishes to select. As such, increased accuracy during ambiguous state trials is explainable by the statistics of the task. Given their success and the equal opportunity that female and male mice have to utilize ambiguous states, it remains unclear why females are more likely to occupy ambiguous states, while the model finds that males remain in exploration longer. One possible explanation is that the neural representation of rule states could be more defined in females. A similar ambiguous “transition-like” state has been observed in neural data from the medial prefrontal cortex in some, but not all, male rats during a rule shifting task (Durstewitz et al., 2010), and the establishment of neural representations in frontal cortex is needed for exploiting a rule (Ebitz et al., 2020). Future work may benefit from neural recording techniques to further understand the origin of ambiguous rule states, and to determine if sex differences in cortical representation of rule states accounts for this difference.

In this task, higher cognitive flexibility in females compared to males relies on high response stability in females. While this may seem counterintuitive, we suggest that cognitive flexibility can be formulated as the rate of adaptive transitions between exploration and exploitation, and that our modeling approach captures this and could be useful in measuring cognitive flexibility across other task designs as well. In particular, female mice showed an increased ability to transition from exploration into exploiting rule states as measured by an input-output hidden Markov model, and we believe that this is due to increases in choice bias and learning rate in females as identified by the reinforcement learning drift diffusion model. An increased ability to move from exploration into exploit is a particularly effective strategy for cognitive flexibility tasks that require patterns of sustained correct performance to detect increased error rates when the correct response has changed to engage cognitive flexibility. Our results suggest that we may be able to quantify cognitive flexibility at a computational level using hidden Markov models by measuring explore-exploit state transitions, which in the future may permit measurements of cognitive flexibility in broader decision making contexts, for example in bandit tasks.

Across species, cognitive flexibility is traditionally measured during rule shifting tasks (Darrah et al., 2008; Reimer et al., 2024). Designed to be homologous to human assays of flexibility, including the Multi Source Interference Test and the Wisconsin Card Sorting Test, Attentional Set Shifting (Set Shift) tasks have been used to reveal neural dynamics important for attending to and shifting between attentional “sets” or rules within a single session in rats and non-human primates (Darrah et al., 2008; Ebitz et al., 2020, 2019; Reimer et al., 2024). While attempts to create mouse Set Shift tasks have been published (Bissonette and Powell, 2012; Heisler et al., 2015; Parikh et al., 2016), the field has lacked an operant rule shifting task for mice that permits continuous rule shifts within a single session. Having a Set Shift task adapted for mice that permits temporal sensitivity has long been a goal in the field (Bissonette and Powell, 2012) given the genetic tractability of mouse models and our ability to measure neural dynamics relies on within-session measures. Unlike previously published set shifting tasks in mice, our current approach both allows us to ask mice to shift between rules repeatedly within a session while also permitting the high throughput and controllability of operant testing. An operant design provides a well defined trial structure, a repeatable within-subjects design, and multiple within-session rule shifts with no investigator intervention required. These features make our Set Shift task ideal for addressing neurophysiological questions, as well as asking about sources of individual differences via genetic, viral, and circuit manipulations. The robust sex differences we have observed point to potential neural mechanisms that may drive differences in this critical executive function that can be explored in future research.

## Acknowledgements

This work was supported by the US National Institutes of Health: R01NS120851 (ASW), R01MH124687 (ASW), R01MH123661 (NMG), P50MH119569 (NMG), T32NS105604 (NG), and by Canada Research Chair CRC-2022-00192 (RBE) and NSERC RGPIN-2020-05577 (RBE). ASW acknowledges support from the MnDRIVE Brain Conditions Initiative and the Minnesota Medical Discovery Team on Addictions. We additionally acknowledge the Minnesota Supercomputing Institute (MSI) at the University of Minnesota for providing resources that contributed to the research results reported within this paper (URL: http://www.msi.umn.edu). We thank Dana Mueller, Elizabeth Sachse, A Yang, Laura Garbe, and Kira Stetler for technical assistance.

## Supplemental Figures

**Supp. Figure 1.**
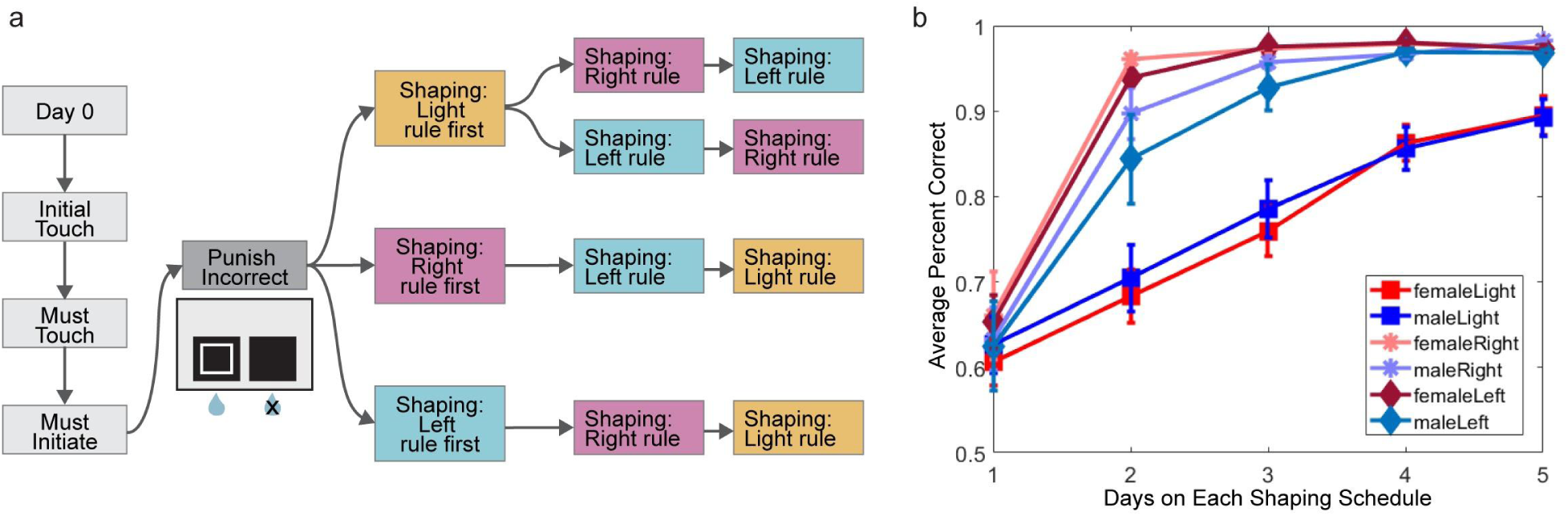
(a) Schematic of the training schedules that mice complete prior to testing on the Set Shift task. Schedules in light grey (Day 0, Initial Touch, Must Touch, and Must Initiate) have been previously described in Chen et al., 2021 and are used to train basic touchscreen use. For the Set Shift training pipeline, Punish Incorrect has been modified to reinforce that the touchscreen is active when the outlined white square is present on the screen. After Punish Incorrect, animals learned each rule individually and were counterbalanced across shaping schedules. (b) By the end of the fifth day on each shaping schedule, all animals, regardless of sex or training order, learned each rule as measured by average percent correct. By day five on each shaping schedule, sex differences in performance of each individual rule stabilized. After learning, performance on the Side Rule shaping schedules was significantly higher compared to Light Rule shaping on day five regardless which rule dimension animals learned first (p=8.8956e-8).

**Supp. Figure 2.**
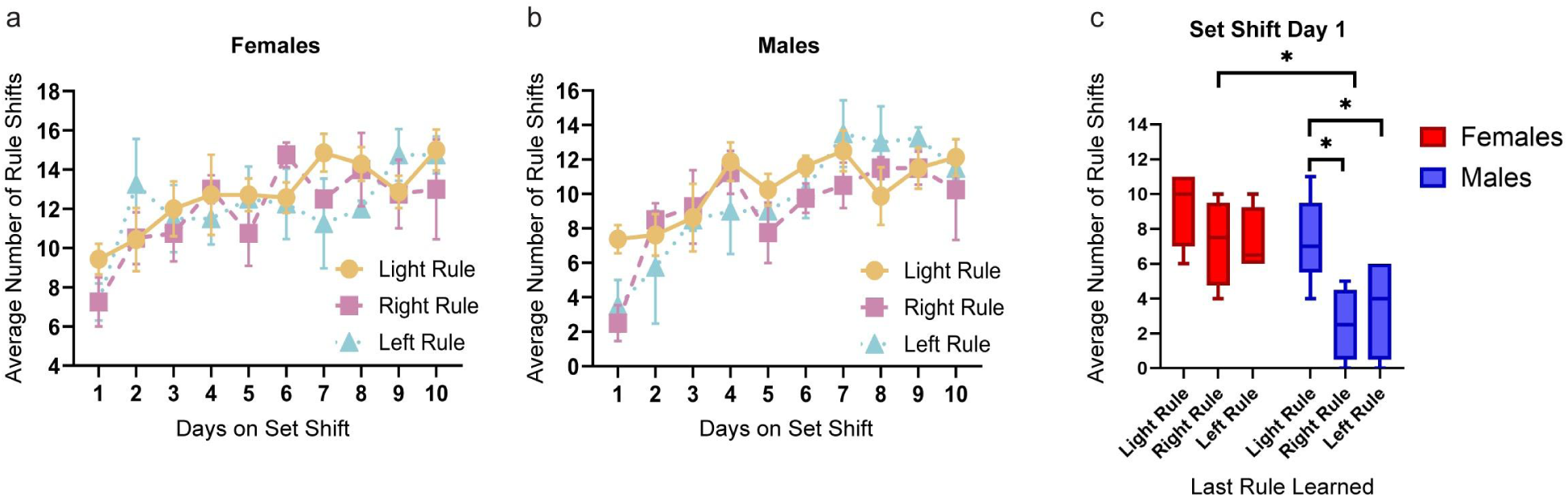
Prior to the first day on the full Set Shift task (Days on Set Shift Day 1), animals completed individual rule shaping where each rule was learned individually. Training order groups in this figure represent the rule shaping that animals completed last, immediately preceding testing on the full Set Shift task. (a) Average number of rule shifts that females in each training group (light shaping last (n=7), right shaping last (n=4), and left shaping last (n=4)) completed throughout the 10 days on the Set Shift task. (b) Average number of rule shifts that males in each training group (light shaping last (n=8), right shaping last (n=4), and left shaping last (n=4)) completed throughout the 10 days on the Set Shift task. (c) Overall, the average number of rule shifts completed during Day 1 on Set Shift was influenced by the last shaping rule that animals completed (p=0.0022) and sex (p=0.0004). Directly comparing performance of each group (sex by training order) revealed that between groups, female mice that completed right shaping last completed significantly more rule shifts on the first day of Set Shift compared to males that completed right shaping last (p=0.0276). Within sex effects were also found–males that completed side shaping last performed significantly fewer rule shifts on Day 1 of Set Shift compared to males that completed light shaping last (left-light comparison: p=0.0323, right-light comparison: p=0.0055).

**Supp. Figure 3.**
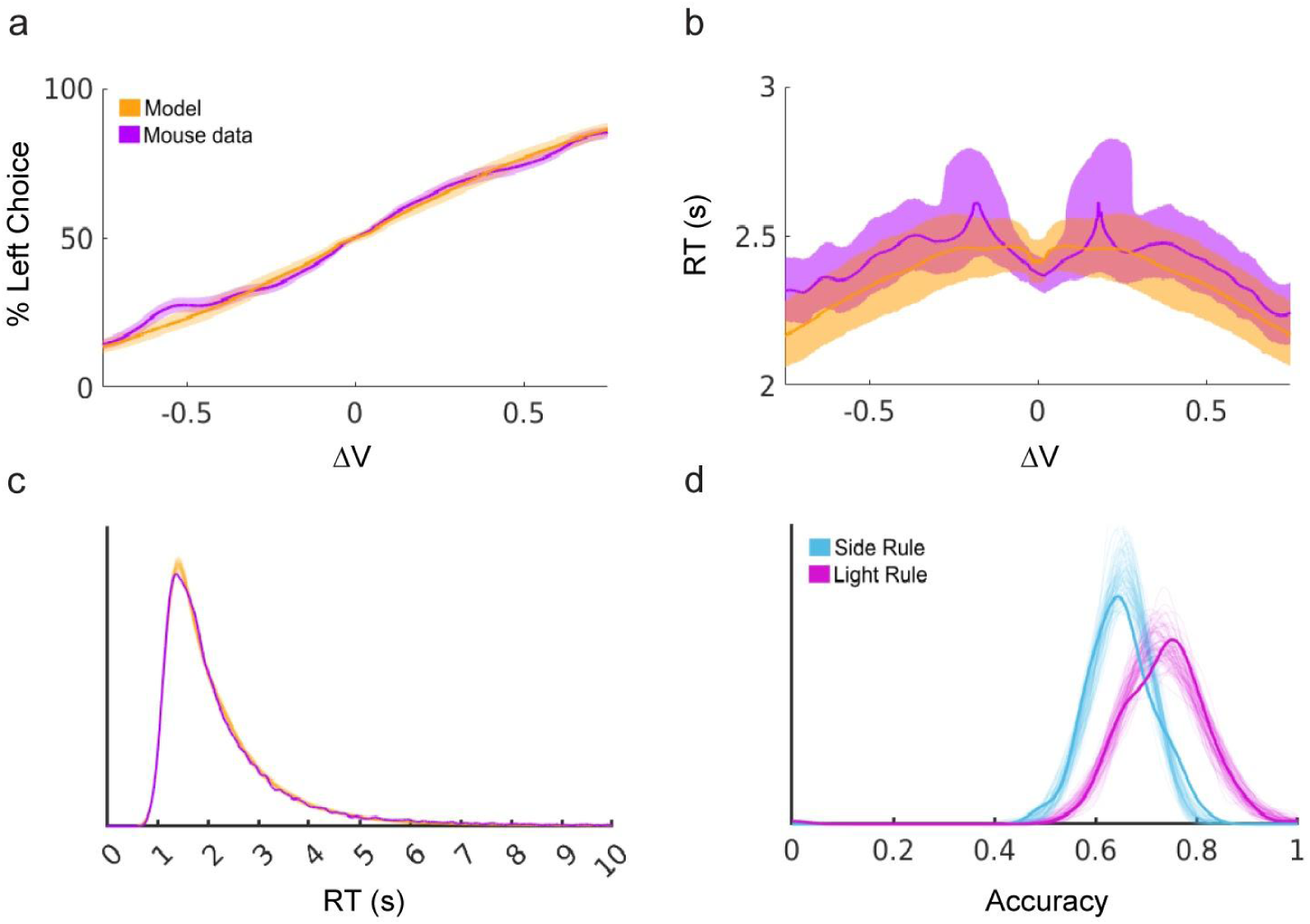
(a) In both the modeled data (orange) and mouse behavioral data (purple), the left choice was selected least when its value was low and most when its value was high. (b) Response times (RTs) (seconds) were fastest when the value between each choice (**Δ**V) differed the most, and slower as the values became more similar in both the mouse data (purple) and model data (orange). (c) Overall, RTs from the model (orange) did not significantly differ from RTs in the mouse data (purple). d) Mouse choice accuracy (transparent) in each rule type (side: blue, light: purple) were similarly captured by the model data (opaque).

**Supp. Figure 4.**
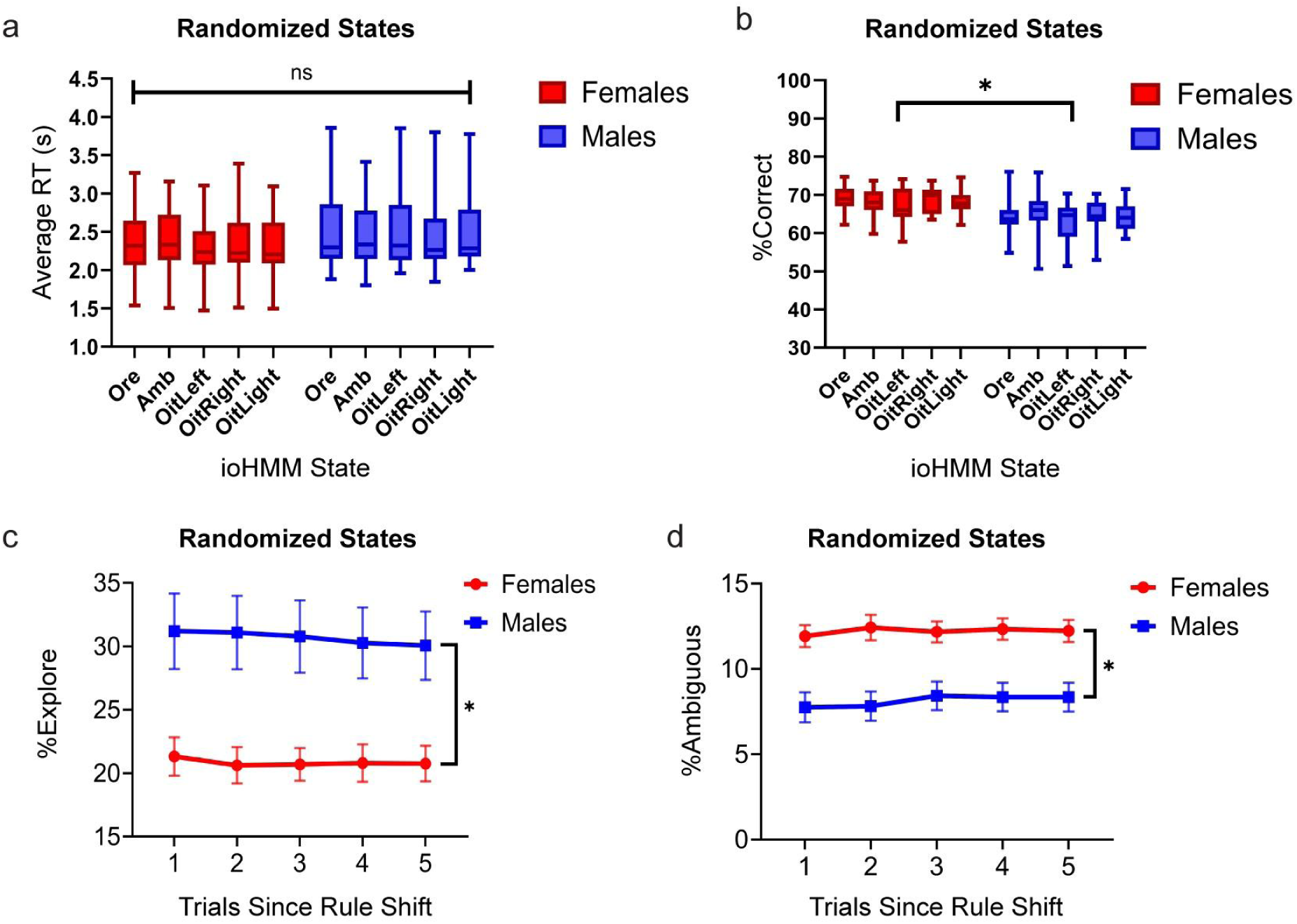
To validate our ioHMM findings, we randomized the trial-by-trial state sequences for each animal, each session of Set Shift. This allowed us to uncouple state labels from their respective trials without changing the overall composition of states for each animal. (a) No effect of state on average response time (RT) when state labels are randomized across trial sequences (p=0.9961). (b) No effect of state on average percent of correct trials when state labels are randomized (p=0.5025). The overall effect of sex on percent correct remains, with female mice demonstrating higher accuracy compared to males (p=3.30226e-07). (c) After randomizing state labels, the effect of trial (temporal structure) is no longer significant on the percent of explore labeled trials 1-5 trials following rule shifts (p=0.9969). The overall effect of sex on exploration remains, with increased exploration in male mice (p=2.81896e-10). (d) The effect of trial on percent of ambiguous labeled states (trials with two equiprobable exploit states) is no longer significant when state labels are randomized across trials (p=0.9636). Overall, the number of ambiguous states in each sex remains unchanged, with significantly more ambiguous states occurring in female mice (p=3.44438e-14).

**Supp. Figure 5.**
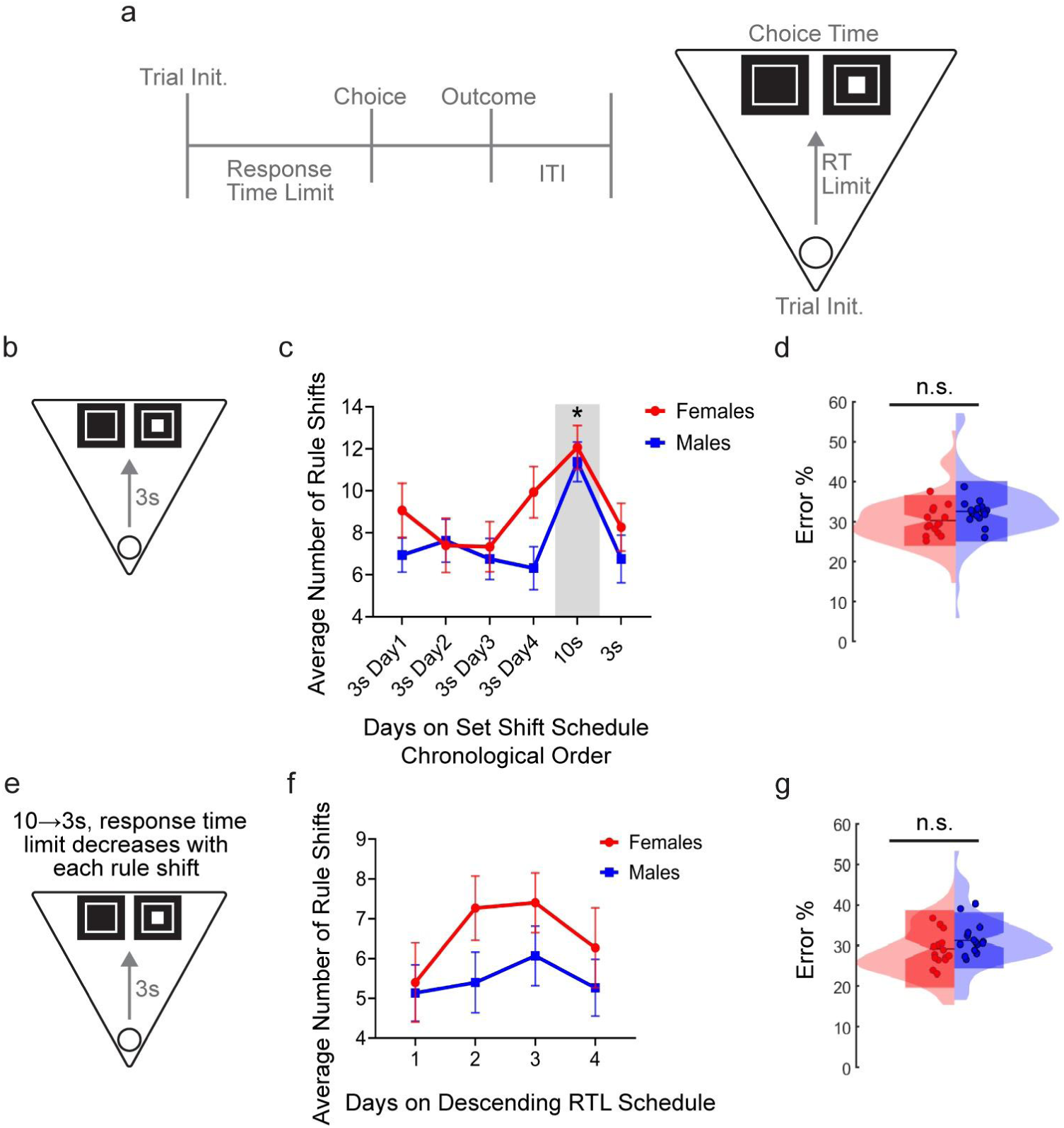
(a) Schematic representation of the trial structure for all Set Shift schedules. After trial initiation, the animal has 10-3 seconds to make a choice (response time limit) depending on schedule. The response time limit (RT Limit) is represented in the top-down chamber illustration by an arrow from trial initiation at the magazine to choice time at the touchscreen. (b) Illustration of the Set Shift schedule with a three second response time limit. (c) The average number of rule shifts that female (red) and male (blue) mice completed during each chronological day on the three second Set Shift schedule. After four days on the three second schedule, animals were again tested on our standard Set Shift schedule which allowed a 10 second response time limit. The average number of completed rule shifts significantly improved during the 10 second Set Shift schedule (p=0.0003) and again fell when tested on the three second Set Shift schedule the following day. Overall, testing on a schedule with a three second response time limit minimized sex differences in the average number of rule shifts completed. (d) The effect of sex on the percent of errors performed during Set Shift is no longer significant during the three second Set Shift schedule (p=0.40762). (e) Illustration of the descending Set Shift schedule where the response time limit starts at 10 seconds and decreases by one second with each completed rule shift until the response time limit reaches three seconds. Data from this schedule in f and g are specifically from trials at the three second response time limit. (f) No significant effect of sex or day on the average number of rule shifts completed at the three second response time limit in this schedule (p=0.0561). (g) No significant sex difference in accuracy (Error %) during trials at the three second response time limit within this schedule (p=0.5312).

## References

Addicott MA, Pearson JM, Sweitzer MM, Barack DL, Platt ML. 2017. A primer on foraging and the explore/exploit trade-off for psychiatry research. Neuropsychopharmacology 42:1931–1939.

Basu I, Yousefi A, Crocker B, Zelmann R, Paulk AC, Peled N, Ellard KK, Weisholtz DS, Cosgrove GR, Deckersbach T, Eden UT, Eskandar EN, Dougherty DD, Cash SS, Widge AS. 2023. Closed-loop enhancement and neural decoding of cognitive control in humans. Nat Biomed Eng 7:576–588.

Bengio Y, Frasconi P. 1994. An Input Output HMM Architecture In: Tesauro G, Touretzky D, Leen T, editors. Advances in Neural Information Processing Systems. MIT Press.

Bissonette GB, Powell EM. 2012. Reversal learning and attentional set-shifting in mice. Neuropharmacology 62:1168–1174.

Blimes JA. 1998. A gentle tutorial of the EM algorithm and its application to parameter estimation for Gaussian mixture and hidden Markov models. Int Comput Sci Inst 4.

Burguière E, Monteiro P, Mallet L, Feng G, Graybiel AM. 2015. Striatal circuits, habits, and implications for obsessive-compulsive disorder. Curr Opin Neurobiol 30:59–65.

Chen CS, Ebitz RB, Bindas SR, Redish AD, Hayden BY, Grissom NM. 2021a. Divergent Strategies for Learning in Males and Females. Curr Biol 31:39–50.e4.

Chen CS, Knep E, Han A, Ebitz RB, Grissom NM. 2021b. Sex differences in learning from exploration. Elife 10. doi:10.7554/eLife.69748

Cox J, Minerva AR, Fleming WT, Zimmerman CA, Hayes C, Zorowitz S, Bandi A, Ornelas S, McMannon B, Parker NF, Witten IB. 2023. A neural substrate of sex-dependent modulation of motivation. Nat Neurosci 26:274–284.

Cuthbert BN, Insel TR. 2013. Toward the future of psychiatric diagnosis: the seven pillars of RDoC. BMC Med 11:126.

Darrah JM, Stefani MR, Moghaddam B. 2008. Interaction of N-methyl-D-aspartate and group 5 metabotropic glutamate receptors on behavioral flexibility using a novel operant set-shift paradigm. Behav Pharmacol 19:225–234.

de Oliveira AR, Reimer AE, Simandl GJ, Nagrale SS, Widge AS. 2021. Lost in translation: no effect of repeated optogenetic cortico-striatal stimulation on compulsivity in rats. Transl Psychiatry 11:315.

Durstewitz D, Huys QJM, Koppe G. 2021. Psychiatric illnesses as disorders of network dynamics. Biol Psychiatry Cogn Neurosci Neuroimaging 6:865–876.

Durstewitz D, Vittoz NM, Floresco SB, Seamans JK. 2010. Abrupt transitions between prefrontal neural ensemble states accompany behavioral transitions during rule learning. Neuron 66:438–448.

Ebitz RB, Albarran E, Moore T. 2018. Exploration Disrupts Choice-Predictive Signals and Alters Dynamics in Prefrontal Cortex. Neuron 97:450–461.e9.

Ebitz RB, Sleezer BJ, Jedema HP, Bradberry CW, Hayden BY. 2019. Tonic exploration governs both flexibility and lapses. PLoS Comput Biol 15:e1007475.

Ebitz RB, Tu JC, Hayden BY. 2020. Rules warp feature encoding in decision-making circuits. PLoS Biol 18:e3000951.

Gelman A, Rubin DB. 1992. Inference from iterative simulation using multiple sequences. Stat Sci 7:457–472.

Golden CEM, Martin AC, Kaur D, Mah A, Levy DH, Yamaguchi T, Lasek AW, Lin D, Aoki C, Constantinople CM. 2023. Estrogenic control of reward prediction errors and reinforcement learning. bioRxiv. doi:10.1101/2023.12.09.570945

Grissom NM, Glewwe N, Chen C, Giglio E. 2024. Sex mechanisms as nonbinary influences on cognitive diversity. Horm Behav 162:105544.

Grissom NM, Herdt CT, Desilets J, Lidsky-Everson J, Reyes TM. 2015. Dissociable deficits of executive function caused by gestational adversity are linked to specific transcriptional changes in the prefrontal cortex. Neuropsychopharmacology 40:1353–1363.

Grissom NM, Reyes TM. 2019. Let’s call the whole thing off: evaluating gender and sex differences in executive function. Neuropsychopharmacology 44:86–96.

Heisler JM, Morales J, Donegan JJ, Jett JD, Redus L, O’Connor JC. 2015. The attentional set shifting task: a measure of cognitive flexibility in mice. J Vis Exp. doi:10.3791/51944

Horner AE, Heath CJ, Hvoslef-Eide M, Kent BA, Kim CH, Nilsson SRO, Alsiö J, Oomen CA, Holmes A, Saksida LM, Bussey TJ. 2013. The touchscreen operant platform for testing learning and memory in rats and mice. Nat Protoc 8:1961–1984.

Levy DR, Hunter N, Lin S, Robinson EM, Gillis W, Conlin EB, Anyoha R, Shansky RM, Datta SR. 2023. Mouse spontaneous behavior reflects individual variation rather than estrous state. Curr Biol 33:1358–1364.e4.

Makowski D, Ben-Shachar MS, Chen SHA, Lüdecke D. 2019. Indices of effect existence and significance in the Bayesian framework. Front Psychol 10:2767.

Mar AC, Horner AE, Nilsson SRO, Alsiö J, Kent BA, Kim CH, Holmes A, Saksida LM, Bussey TJ. 2013. The touchscreen operant platform for assessing executive function in rats and mice. Nat Protoc 8:1985–2005.

Merikangas AK, Almasy L. 2020. Using the tools of genetic epidemiology to understand sex differences in neuropsychiatric disorders. Genes Brain Behav 19:e12660.

Murphy KP. 2021. Machine Learning. MIT Press. https://mitpress.mit.edu/9780262018029/machine-learning/

Orsini CA, Brown TE, Hodges TE, Alonso-Caraballo Y, Winstanley CA, Becker JB. 2022. Neural mechanisms mediating sex differences in motivation for reward: Cognitive bias, food, gambling, and drugs of abuse. J Neurosci 42:8477–8487.

Orsini CA, Setlow B. 2017. Sex differences in animal models of decision making. J Neurosci Res 95:260–269.

Palmer JA, White SR, Chavez Lopez K, Laubach M. 2024. The role of the rat prefrontal cortex and sex differences in decision-making. J Neurosci 44:e0550242024.

Parikh V, Naughton SX, Yegla B, Guzman DM. 2016. Impact of partial dopamine depletion on cognitive flexibility in BDNF heterozygous mice. Psychopharmacology (Berl) 233:1361–1375.

Rabiner LR. 1989. A tutorial on hidden Markov models and selected applications in speech recognition. Proc IEEE Inst Electr Electron Eng 77:257–286.

Reimer AE, Dastin-van Rijn EM, Kim J, Mensinger ME, Sachse EM, Wald A, Hoskins E, Singh K, Alpers A, Cooper D, Lo M-C, de Oliveira AR, Simandl G, Stephenson N, Widge AS. 2024. Striatal stimulation enhances cognitive control and evidence processing in rodents and humans. Sci Transl Med 16. doi:10.1126/scitranslmed.adp1723

Uddin LQ. 2021. Cognitive and behavioural flexibility: neural mechanisms and clinical considerations. Nat Rev Neurosci 22:167–179.

Widge AS, Zorowitz S, Basu I, Paulk AC, Cash SS, Eskandar EN, Deckersbach T, Miller EK, Dougherty DD. 2019. Deep brain stimulation of the internal capsule enhances human cognitive control and prefrontal cortex function. Nat Commun 10:1536.

Wiecki TV, Sofer I, Frank MJ. 2013. HDDM: Hierarchical Bayesian estimation of the Drift-Diffusion Model in Python. Front Neuroinform 7:14.

Yoest KE, Quigley JA, Becker JB. 2018. Rapid effects of ovarian hormones in dorsal striatum and nucleus accumbens. Horm Behav 104:119–129.

